# Nonlinear Mixed-Effects and Full Bayesian Population Pharmacokinetic Analysis of Ceftolozane–Tazobactam in Critically Ill Patients

**DOI:** 10.64898/2026.03.24.713879

**Authors:** Paulina M. Okuńska, Michał Borys, Elżbieta Rypulak, Paweł Piwowarczyk, Marta Szczukocka, Grzegorz Raszewski, Mirosław Czuczwar, Paweł Wiczling

## Abstract

Pharmacokinetic studies in critically ill patients are often constrained by small sample sizes, limiting the strength and generalizability of conclusions drawn solely from observed data. Bayesian inference offers a powerful strategy to address this challenge by incorporating prior knowledge. In this study, we evaluated two model-based approaches for characterizing the population pharmacokinetics of ceftolozane and tazobactam in critically ill patients, comparing nonlinear mixed-effects modeling with Bayesian hierarchical analyses. The Bayesian methods incorporated literature-derived prior information. The data was collected from 13 critically ill patients receiving 3.0 g of ceftolozane combined with tazobactam (2:1) via intravenous infusion. Pharmacokinetic modeling was performed using NONMEM and Stan software with the Torsten extension. Model diagnostics and graphical analyses were conducted in RStudio with relevant packages. In the absence of prior information, a one-compartment model with a limited set of parameters describing inter-individual variability adequately characterized the pharmacokinetics of ceftolozane and tazobactam. When prior information was incorporated, a two-compartment model became feasible and yielded a characterization of parameter variability and correlations that was more consistent with published literature. The application of Bayesian inference ensured alignment with existing literature on ceftolozane and tazobactam pharmacokinetics and mitigated some systematic biases observed in the data-driven approaches. Moreover, the Bayesian approach enables direct decision-making by incorporating uncertainty into the analysis, as demonstrated by probability of target attainment analysis. Collectively, these results underscore the utility of Bayesian methods in pharmacokinetic modeling for critically ill patients, offering a robust framework for optimizing dosing strategies in data-limited settings.

## 2. Introduction

The increasing prevalence of antibiotic resistance underscores the critical importance of research on antibiotics, particularly in the context of improving dosing regimens for bacterial infections. Administering an insufficient dose over an inadequate duration may result in incomplete treatment of the disease, ultimately leading to bacterial resistance to the antibiotic or its class. Conversely, an excessive dose or prolonged therapy can lead to adverse effects. Within the general patient population, certain subgroups exhibit altered pharmacokinetics that may require dose adjustments. These include, among others, pediatric patients, oncology patients, and critically ill individuals. Conducting pharmacokinetic studies in these subpopulations can be challenging due to difficulties in patient recruitment. As a result, such studies often include only small cohorts of participants, which limits the ability to draw robust conclusions [1,2].

Nonlinear mixed-effects (NLME) modeling is the most used method for analyzing clinical pharmacokinetic data [3]. This hierarchical approach includes two types of parameters: population-level (fixed effects) and individual-level (random effects). Fixed effects describe characteristics assumed to be consistent across the entire population, whereas random effects represent parameters that vary between individuals, capturing inter-individual variability. These models enable identification of predictive covariates and comparison of variability between the overall population and specific subgroups. NLME models estimate fixed and random effects using likelihood-based methods, producing point estimates with uncertainty derived through approximation techniques [4]. To strengthen pharmacokinetic analyses, particularly when data are limited, prior knowledge from literature or other studies can be incorporated using Bayesian methods. In a full Bayesian framework, all parameters (including random effects) are treated probabilistically, prior information is incorporated explicitly, and the model yields full posterior distributions rather than single point estimates, providing a transparent quantification of how new data refines prior assumptions [5].

In 2015, the European Medicines Agency (EMA) approved the marketing of the novel drug ceftolozane/tazobactam, which combines a β-lactam antibiotic (a cephalosporin) with a β-lactamase inhibitor [6,7]. It was recommended for the treatment of complicated intra-abdominal infections, severe kidney infections, and complicated urinary tract infections, as well as hospital-acquired pneumonia, including ventilator-associated pneumonia, in adult patients. Since 2022, these indications have been extended to the pediatric population for the first two indications [8]

This antibiotic is particularly important, as it finds specific application in treating infections caused by multidrug-resistant (MDR) *Pseudomonas aeruginosa* and extended-spectrum β-lactamase (ESBL)-producing *Enterobacterales*, which are challenging to manage due to their extensive resistance mechanisms against previously known antibiotics. Therefore, research on this drug and understanding the need for dosing optimization in subpopulations may enhance our chances of success in treating severe bacterial infections [9,10].

## 3. Methods

### 3.1 Study Design

Patients enrolled in the study were treated at the Second Department of Anesthesiology and Intensive Care of the Independent Public Clinical Hospital No. 1 in Lublin. Following approval from the Bioethics Committee (approval number: KE-0254/258/2014), 13 patients were qualified for the study, in whom the following diseases were diagnosed: viral pneumonia, pneumonia with acute respiratory distress syndrome (ARDS), acute pancreatitis, perforated gastric ulcer, abdominal aortic aneurysm with stent-graft replacement by a prosthesis, cardia cancer combined with pneumonia, gastric cancer, uterine fibroid with evisceration, and ovarian cancer.

In two patients, due to severe pneumonia, acute respiratory distress syndrome, and multi-organ failure, continuous extracorporeal membrane oxygenation (ECMO) was implemented. In five patients, continuous veno-venous hemofiltration (CRRT) was applied, with dialysis therapy initiated on the second day of antibiotic treatment in two of them, and one of whom died.

### 3.2 Treatment Procedure

Patients were qualified for antibiotic therapy with Zerbaxa® (Merck Sharp & Dohme B.V., Netherlands) due to the necessity of initiating empirical treatment. The antibiotic was administered intravenously at a dose of 3.0 g (2.0 g ceftolozane + 1.0 g tazobactam) q8h, which constitutes the standard dosing regimen for the treatment of hospital-acquired pulmonary infections as specified by the manufacturer [11]. The infusion duration was 3 hours in 11 patients, with one dose in one patient administered as a 1-hour infusion. In the case of the two remaining patients, who were treated with ECMO support, the drug administration time was shortened to 1 hour. Notably, a 1-hour infusion is recommended in the standard dosing regimen.

Ceftolozane with tazobactam was prepared immediately prior to administration to patients. For reconstitution, 50 mL of 0.9% sodium chloride solution (Natrium Chloratum, Fresenius Kabi, Poland) was used as the solvent. The resulting drug was administered via continuous infusion using a syringe pump (Perfusor^®^ Space, B Braun, Germany) and a 50-mL syringe (Margomed, Poland). It was delivered through a central venous catheter into the patients’ central vessels, which were the subclavian vein or the internal jugular vein. Table 1 summarizes the patient demographics and clinical characteristics.

**Table 1.**
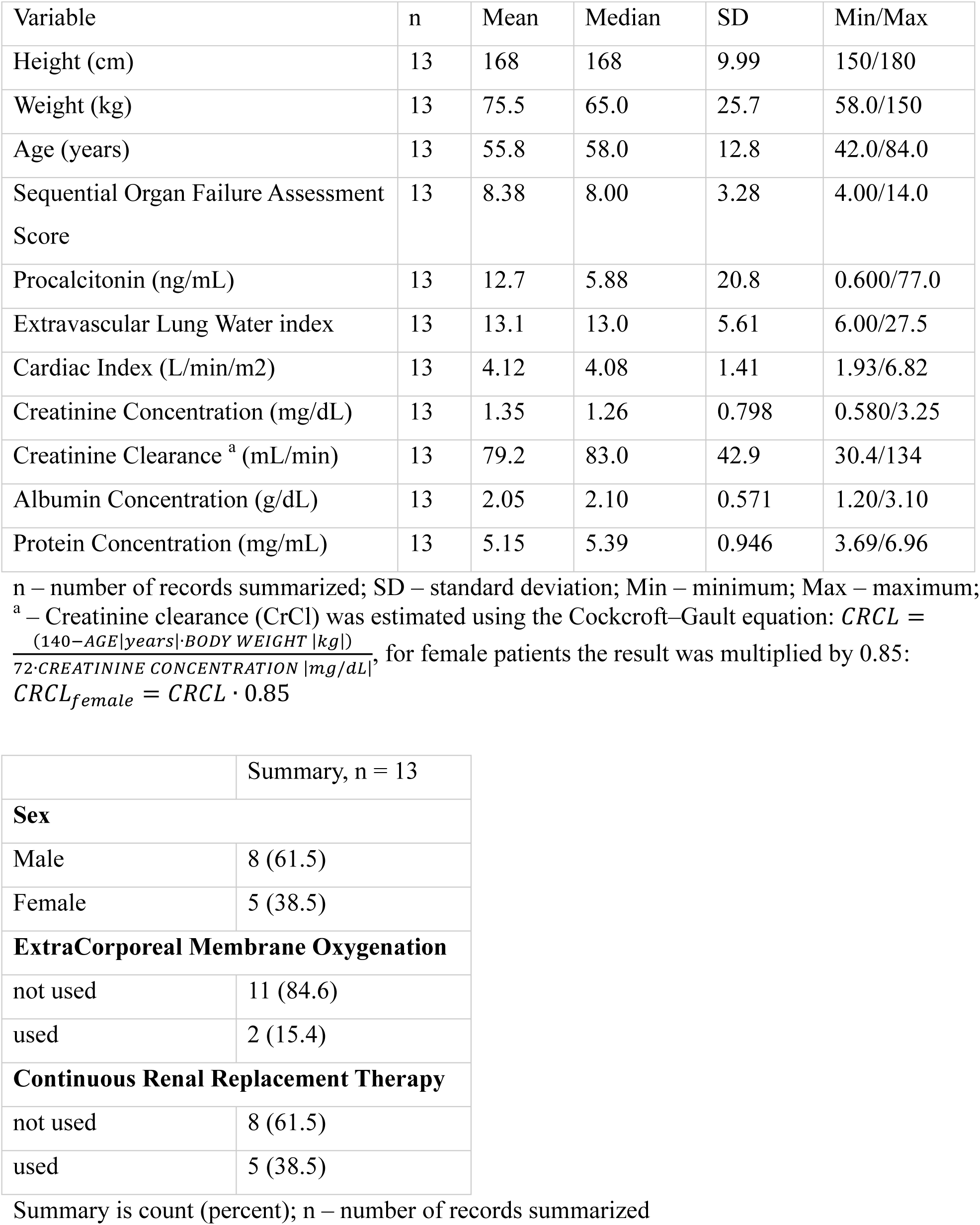
Patient characteristics of the studied population at inclusion.

| Variable | n | Mean | Median | SD | Min/Max |
| --- | --- | --- | --- | --- | --- |
| Height (cm) | 13 | 168 | 168 | 9.99 | 150/180 |
| Weight (kg) | 13 | 75.5 | 65.0 | 25.7 | 58.0/150 |
| Age (years) | 13 | 55.8 | 58.0 | 12.8 | 42.0/84.0 |
| Sequential Organ Failure Assessment Score | 13 | 8.38 | 8.00 | 3.28 | 4.00/14.0 |
| Procalcitonin (ng/mL) | 13 | 12.7 | 5.88 | 20.8 | 0.600/77.0 |
| Extravascular Lung Water index | 13 | 13.1 | 13.0 | 5.61 | 6.00/27.5 |
| Cardiac Index (L/min/m<sup>2</sup>) | 13 | 4.12 | 4.08 | 1.41 | 1.93/6.82 |
| Creatinine Concentration (mg/dL) | 13 | 1.35 | 1.26 | 0.798 | 0.580/3.25 |
| Creatinine Clearance <sup>a</sup> (mL/min) | 13 | 79.2 | 83.0 | 42.9 | 30.4/134 |
| Albumin Concentration (g/dL) | 13 | 2.05 | 2.10 | 0.571 | 1.20/3.10 |
| Protein Concentration (mg/mL) | 13 | 5.15 | 5.39 | 0.946 | 3.69/6.96 |

|  |  |
| --- | --- |
|  | Summary, n = 13 |
| **Sex** |  |
| Male | 8 (61.5) |
| Female | 5 (38.5) |
| **ExtraCorporeal Membrane Oxygenation** |  |
| not used | 11 (84.6) |
| used | 2 (15.4) |
| **Continuous Renal Replacement Therapy** |  |
| not used | 8 (61.5) |
| used | 5 (38.5) |

Summary is count (percent); n – number of records summarized

### 3.3 Plasma Sample Collection and Preparation

During therapy, blood samples were collected to determine drug concentrations. Although the sampling schedule was irregular, most samples were obtained at 3, 5.5, 8, 11, 13.5, 16, 19, and 21.5 h after the first dose, that correspond to samples prior to dosing, at the end of infusion, and 2.5 h after infusion completion. Blood samples were centrifuged immediately after collection, and the resulting plasma was stored in a freezer at − 80 °C.

### 3.4 Bioanalytical Method

Ceftolozane and tazobactam concentrations were quantified using high-performance liquid chromatography (HPLC). Analyses were performed on a Dionex chromatographic system (P580 pump, degasser, mixer, and UVD 170/S/340S variable-wavelength detector) controlled via CHROMELEON™ v.6.30 software with the SummiTox module. Separation was achieved on a Zorbax SB-C18 column (150×4.6 mm, 5 µm). The mobile phase consisted of phosphate buffer (pH 3.0; Fluka) and acetonitrile (15:85, v/v), delivered at a flow rate of 1.0 mL/min. Detection was performed at 215 nm, at which ceftolozane and tazobactam appeared as two distinct peaks. The total run time was 20 minutes. Samples were thawed at room temperature directly before analysis. Samples were prepared by adding 200 µL of acetonitrile to 200 µL of serum in polypropylene tubes. After shaking for 5 minutes, samples were centrifuged at 12,000 rpm for 10 minutes, and a 20 µL aliquot of the supernatant was injected into the liquid chromatography (LC) system. Quantification was performed using calibration curves over a concentration range of 1.0–200 µg/mL.

### 3.5 Model

#### a) Structural model

According to published studies, the pharmacokinetics of ceftolozane and tazobactam are typically described using one- or, more commonly, two-compartment models:

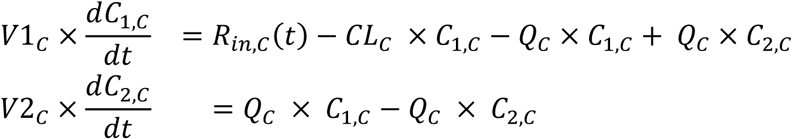

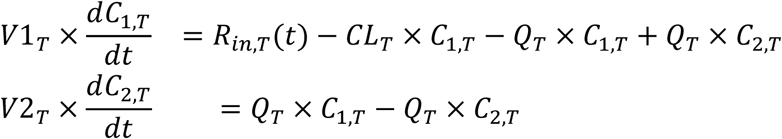

where *R_in_*_,*C*_(*t*) denotes the time-dependent drug administration rate, *C_1_* and *C_2_*represent the concentrations of ceftolozane or tazobactam in the central and peripheral compartments, respectively; *CL* is the clearance, *Q* is the intercompartmental clearance, *V1* and *V2* are the central and peripheral volumes of distribution. The parameters/variables corresponding to ceftolozane or tazobactam are denoted using superscripts (*C* or *T*). In a one-compartment model, *Q* is fixed to 0 and *V2* can be fixed to an arbitrary value.

#### b) Inter-individual variability (IIV)

The pharmacokinetic parameters were modeled using a multivariate log normal distribution:

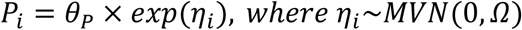

where *θ_P_* is the vector of population (typical) parameters (*CL_C_*, *Q_C_*, *V1_C_*, *V2_C_*, *CL_T_*, *Q_T_*, *V1_T_*, *V2_T_*), *η_i_* are random effects, MVN is a multivariate normal distribution, and *Ω* is the variance–covariance matrix of the random effects. The diagonal elements *diag*(*Ω*) represent the marginal variances, while the off-diagonal elements capture the covariances. In NONMEM we initially assumed covariances for all the parameters were zero and expanded or simplified the random effect model structure based on model diagnostics. For the Bayesian approach, we used a full covariance matrix.

#### c) Observation model

For the residual error model in the NONMEM analysis, we initially applied a proportional error structure:

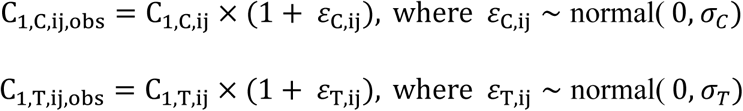

For the Bayesian framework, we instead used an additive model on the log scale with Student-t distributed residuals to improve robustness to outliers:

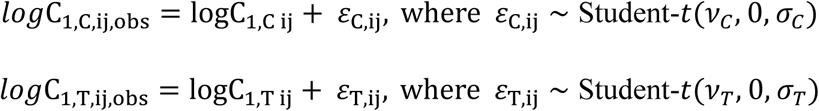

where *C*_1,_*_C_*_,*ij*_ *and C*_1,*T*,*ij*_ are the model-predicted plasma concentrations (for either ceftolozane or tazobactam), *v* is the degrees-of-freedom (normality) parameter, and *σ* is the scale of the corresponding residual error distribution. These error models could be extended to a combined additive and proportional error model based on the model diagnostics.

### 3.6 Nonlinear Mixed-Effects Modeling

The analysis of population pharmacokinetics via nonlinear mixed-effects modeling was performed utilizing the NONMEM program (version 7.3; Icon Development Solutions, USA), along with a Fortran compiler (version 4.6.0) and the bbr package for RStudio [12]. Throughout the entire model development phase, the first-order conditional estimation approach incorporating interaction was applied. Outputs from NONMEM were handled and graphically represented with RStudio® [13].

To compare and select models during development, we evaluated the NONMEM objective function value, standard goodness-of-fit diagnostic plots, and the precision of pharmacokinetic parameter and variability estimates. Model performance was evaluated using Visual Predictive Checks (VPC), calculated from 1,000 datasets simulated with the final parameter estimates [14,15]. To assess parameter uncertainty in the final model, a nonparametric bootstrap procedure was implemented with 500 replicates. From the bootstrap empirical posterior distribution, 90% confidence intervals (5^th^-95^th^ percentile) were obtained for the parameters as described by Parke et al [16]. Further details on the model construction and methods are provided in the Electronic Supplementary Material (ESM).

### 3.7 Bayesian Approach

The Bayesian approach was implemented using Stan/Torsten via the “cmdstanr” [17] and “bbr.bayes” [18] packages in RStudio. Inference used four Markov chains with 1000 iterations each after a 1000-iteration warmup [19]. Convergence was confirmed via Gelman-Rubin statistics and trace plots. R scripts, dataset, and Stan code are available on GitHub. After fitting the model, we examined the MCMC (Markov Chain Monte Carlo) diagnostic suite to verify that the sampling process was proceeding appropriately [20].

Priors were selected to define reasonable ranges for parameters based on pre-data expectations, providing regularization for model stability and incorporating prior evidence to stabilize estimation. Essentially the informative log-normal priors were set to structural parameters from literature [21] with weakly informative priors for IIV and residual error model parameters as follows:

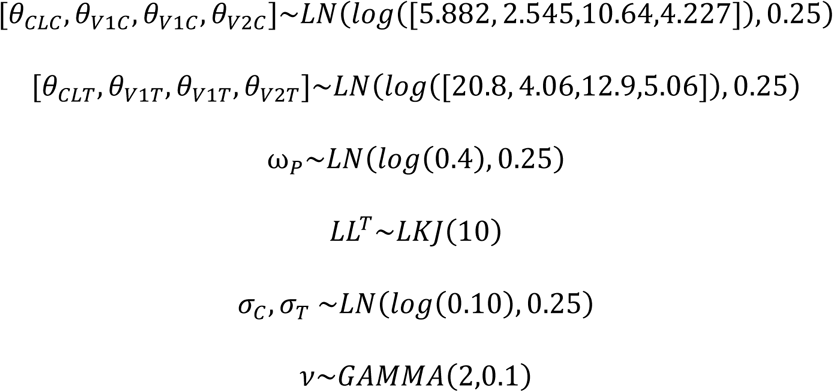

where LN denotes the log-normal distribution, GAMMA denotes the gamma distribution, and LKJ denotes the Lewandowski-Kurowicka-Joe distribution implemented using the Cholesky decompositon. The set of parameters defined within ω*_P_* includes *CL_C_, Q_C_, V1_C_, V2_C_, CL_T_, Q_T_, V1_T_,* and *V2_T_*. For stability the etas were sampled via Cholesky factorization and standard deviations:

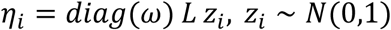

where *L* is the Cholesky factor of the IIV correlation matrix, and z represents the standard normal distribution. As in the case of NMLE analysis, the model performance was evaluated using visual Posterior Predictive Checks (PPC), calculated from 1,000 datasets simulated based on posterior samples.

### 3.8 Probability of Target Attainment

Dose optimization of antibiotics in special populations can be guided by simulations that combine population pharmacokinetic (PK) models with pharmacodynamic (PD) targets to estimate the probability of target attainment (PTA) [22]. For beta-lactam antibiotics, which exhibit time-dependent activity, the key PD index is the percentage of the dosing interval during which free drug concentrations exceed the minimum inhibitory concentration (% fT>MIC). Consequently, increasing drug concentrations above this threshold provides minimal additional benefit on the rate or extent of bacterial killing [23]. PTA analysis used Monte Carlo simulations that incorporated parameter uncertainty and inter-individual variability from the population PK model. Simulated parameter sets were obtained either from the Bayesian posterior distribution or via bootstrap resampling of the NONMEM estimates. Each PTA profile was simulated for 250 individuals across 500 replicates and under various dosing regimens (3g q8h vs. 3g q12h). The PD targets were specified as ≥30% fT > MIC for ceftolozane and ≥20% fT > CT of 1 µg/mL for tazobactam [24]. For each MIC value ranging from 0.25 to 64 mg/L, the proportion of individuals attaining both targets were determined. The proportion for different MIC values was summarized as median (50^th^ percentile) and 5^th^-95^th^ percentile ranges (90% CI) to illustrate the uncertainty in PTA profiles [22]. Simulations for both the recommended q8h regimen and a modified q12h regimen with a 3-hour infusion were conducted using three different simulation approaches: one-compartment model (1CMT COVARIANCE) predictions, bootstrap-based simulations (1CMT BOOTSTRAP) and Bayesian posterior predictive modeling (2CMT BAYES). This allowed us to compare the impact of different analytical approaches on PTA outcomes under a standard and modified dosing regimen. Median PTA values and 5-95% prediction intervals were calculated across MICs.

## 4. Results

### 4.1 Data

The population pharmacokinetic model for ceftolozane co-administered with tazobactam incorporated 101 ceftolozane concentration measurements (8 per day from 12 patients and 5 per day from 1 patient) and 90 tazobactam concentration measurements. Individual patient concentration profiles within this population are illustrated in Figures 1. and S1. Concentrations of both compounds ranged from approximately 1 to 100 μg/mL. The linearity evident in the semi-logarithmic plot suggests that each individual profile could be sufficiently characterized by a one-compartment model; nevertheless, literature predominantly favors a two-compartment model [25–29].

**Figure 1.**
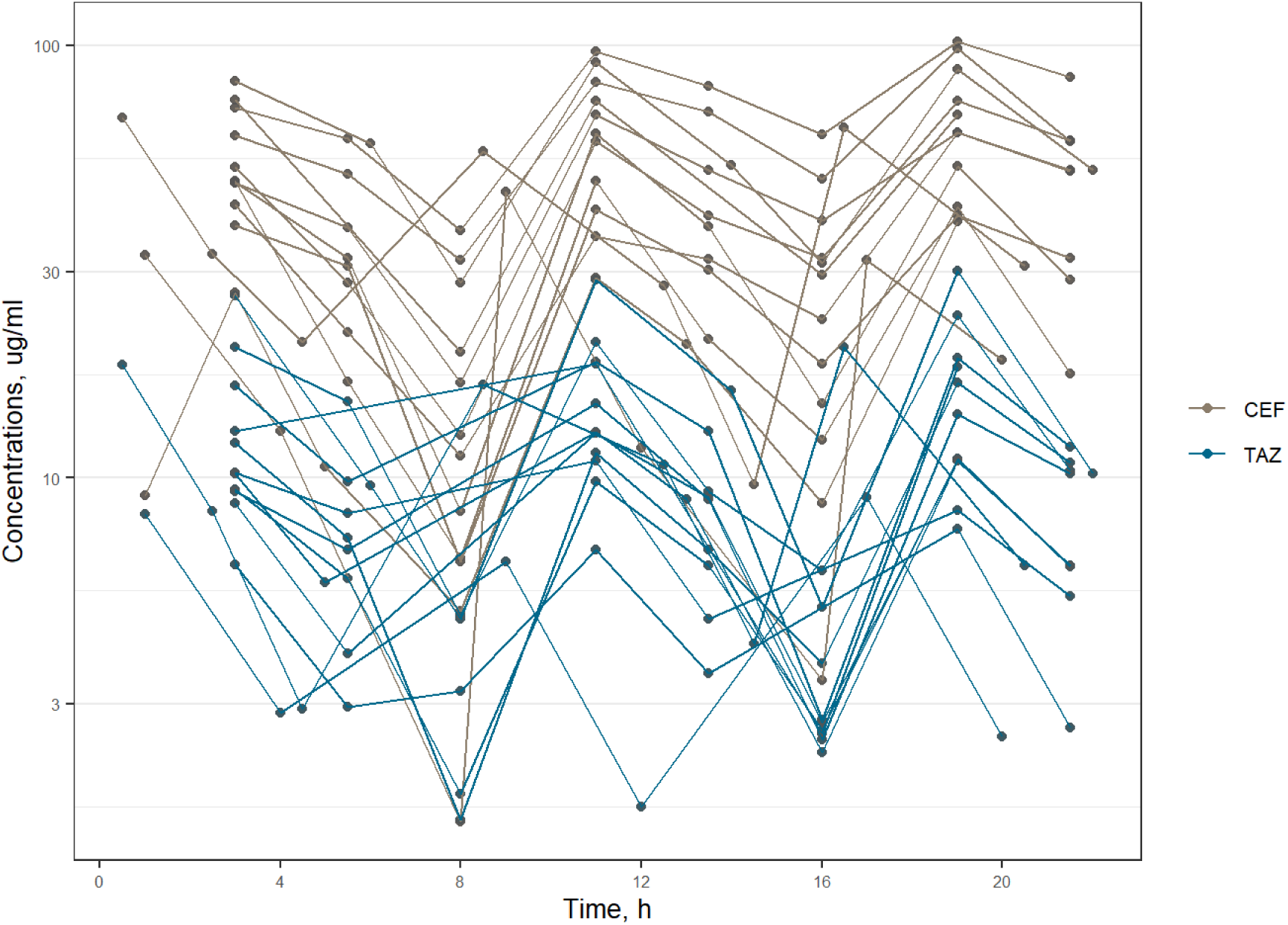
Individual profiles of ceftolozane (CEF, brown) and tazobactam (TAZ, blue) concentrations in plasma on a semi-log scale.

### 4.2 Nonlinear Mixed Effects Modeling

Initially, the available data were analyzed using a standard NMLE workflow using one-and two-compartment models that simultaneously described the pharmacokinetics of ceftolozane and tazobactam. The one-compartment model, with an objective function value (OFV) of 820.5, estimated the parameters of volume of distribution and clearance for both ceftolozane and tazobactam. During model development, individual PK parameters showed significant correlations that were modeled using a full Omega block. Since the correlation between *V_1,T_*and *CL_T_* tended to 1, it was fixed to 1 using shared and scaled ETA. The majority of pharmacokinetic parameters, inter-individual variability, inter-individual covariance, and residual error variances were accurately estimated, with relative standard errors below 50% for most parameters, as presented in Table 2. The two-compartment model yields a higher OFV of 877.838 with parameter estimates presented in Table 2. As the structural model, the simpler one was selected, characterized by a lower OFV and superior goodness-of-fit plots (Figures S3a-b). The inability to estimate the parameters of the two-compartment model may arise from insufficient data or the absence of a second phase in this patient cohort.

**Table 2.**
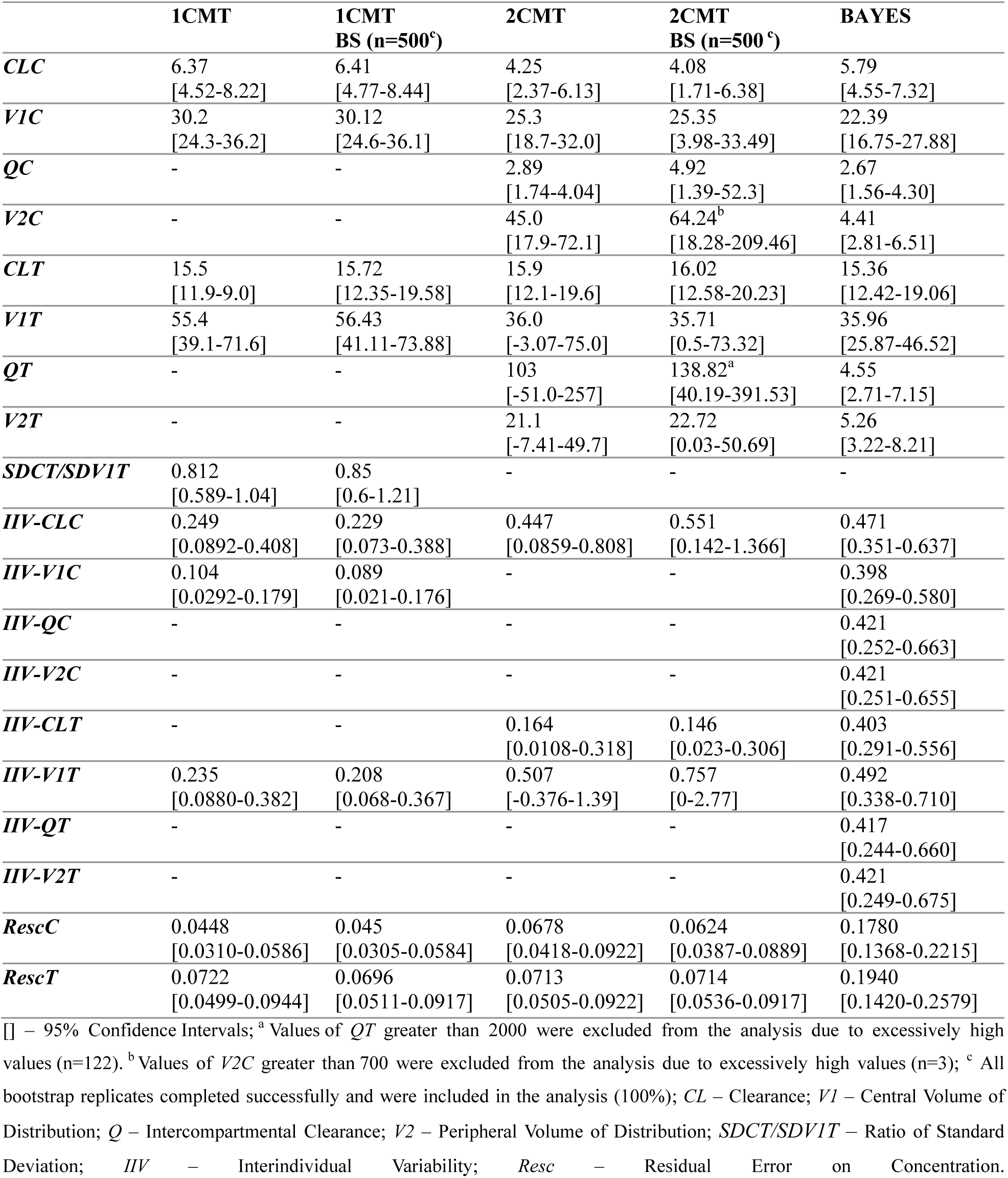
Parameter estimates and standard errors from the one-compartment (1CMT) and two-compartment (2CMT) PK models, their bootstrap analyses, and the Bayesian approach for ceftolozane (C) and tazobactam (T).

The subsequent stage of the analysis involved exploring association between covariates and individual pharmacokinetic parameters. Analysis of the literature and available data suggest the inclusion of the following covariates: age, sex, body weight, creatinine clearance, and presence of CRRT. However, none of the tested variables proved statistically significant, despite evidence from other studies that incorporating creatinine clearance enhances model fitting to the investigated population. It is likely due to a small number of subjects and the relative homogeneity of the study cohort. The comparison of goodness-of-fit plots for both models is presented in Figure 2. In Table S1 there is a comparison of both NONMEM models.

**Figure 2.**
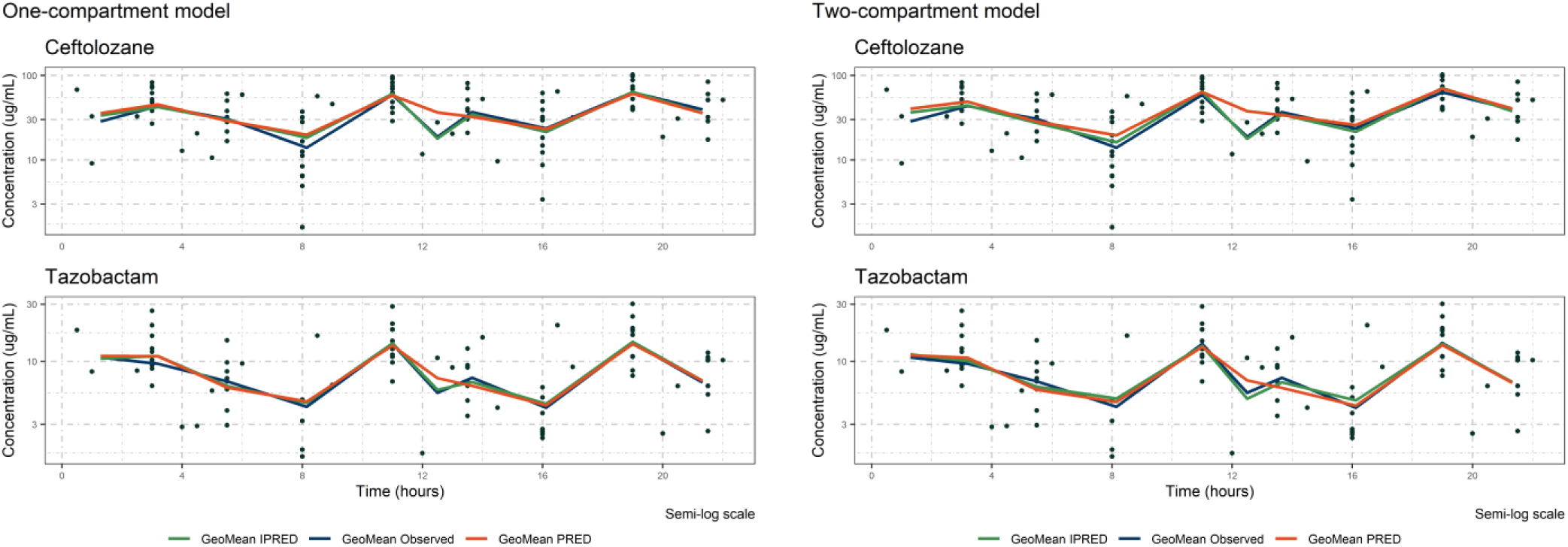
The comparison of goodness-of-fit plots for one-compartment model (left) and two-compartment model (right).

### 4.3 Bayesian Approach

The large number of unknown parameters necessitated model oversimplification under the NMLE approach, potentially leading to unreliable conclusions. To address data sparsity, we therefore employed a full Bayesian approach.

Bayesian population pharmacokinetic models were estimated using Markov Chain Monte Carlo (MCMC) sampling, a computationally intensive approach for approximating complex joint posterior distributions in hierarchical models. Parameter uncertainty was quantified using posterior credible intervals derived from the aggregated MCMC chains, representing the range of parameter values with high posterior probability. MCMC convergence diagnostics are presented in Figures S5a-d. Prior-posterior comparisons for population-level parameters are presented in Figure 3, which illustrates central prior/posterior uncertainty intervals based on quantiles from the prior/posterior draws. The plot displays 50% intervals as thick segments and 90% intervals as thinner outer lines, with points corresponding to the medians.

**Figure 3.**
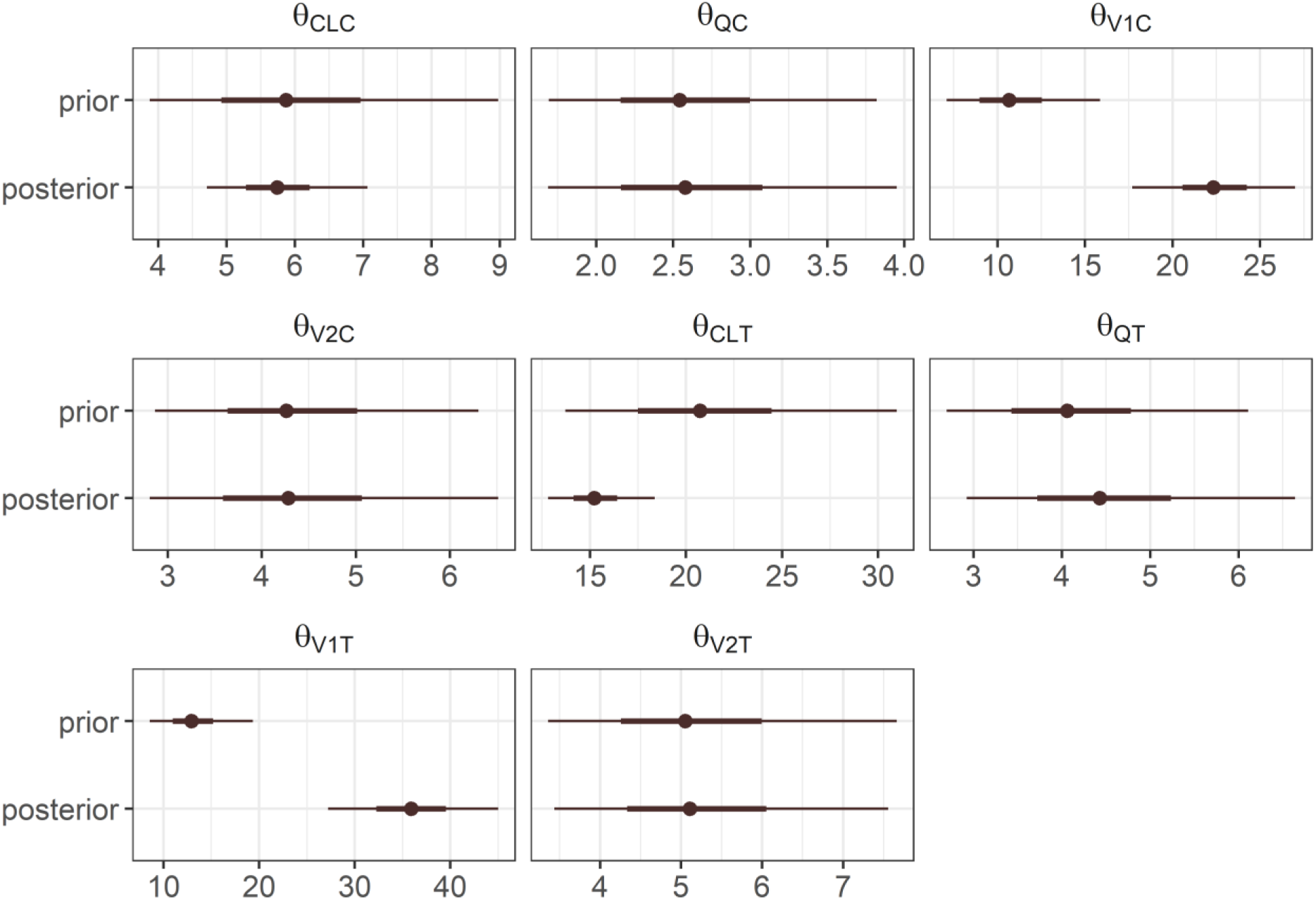
Central posterior uncertainty intervals. The 50% intervals are represented by thick segments, whereas the 90% intervals are depicted by the thinner outer lines. The points illustrated in the plot above correspond to posterior medians.

Model calibration and predictive performance were evaluated using observation-prediction plots (Figure 4). For both analytes, individual posterior predictions aligned closely with the line of identity (*y* = *x*), exhibiting minimal systemic bias and only moderate dispersion at low concentrations (< 1μg/mL). This concordance confirms that the hierarchical Bayesian structure successfully captured interindividual variability in drug exposure. Population predictions showed increased variability, especially at lower concentrations, because they are conditioned only on population-level parameters and do not incorporate individual-specific concentration measurements. Nevertheless, the predictions followed the identity line without pronounced bias, indicating overall adequacy of the two-compartment structural model for both compounds.

**Figure 4.**
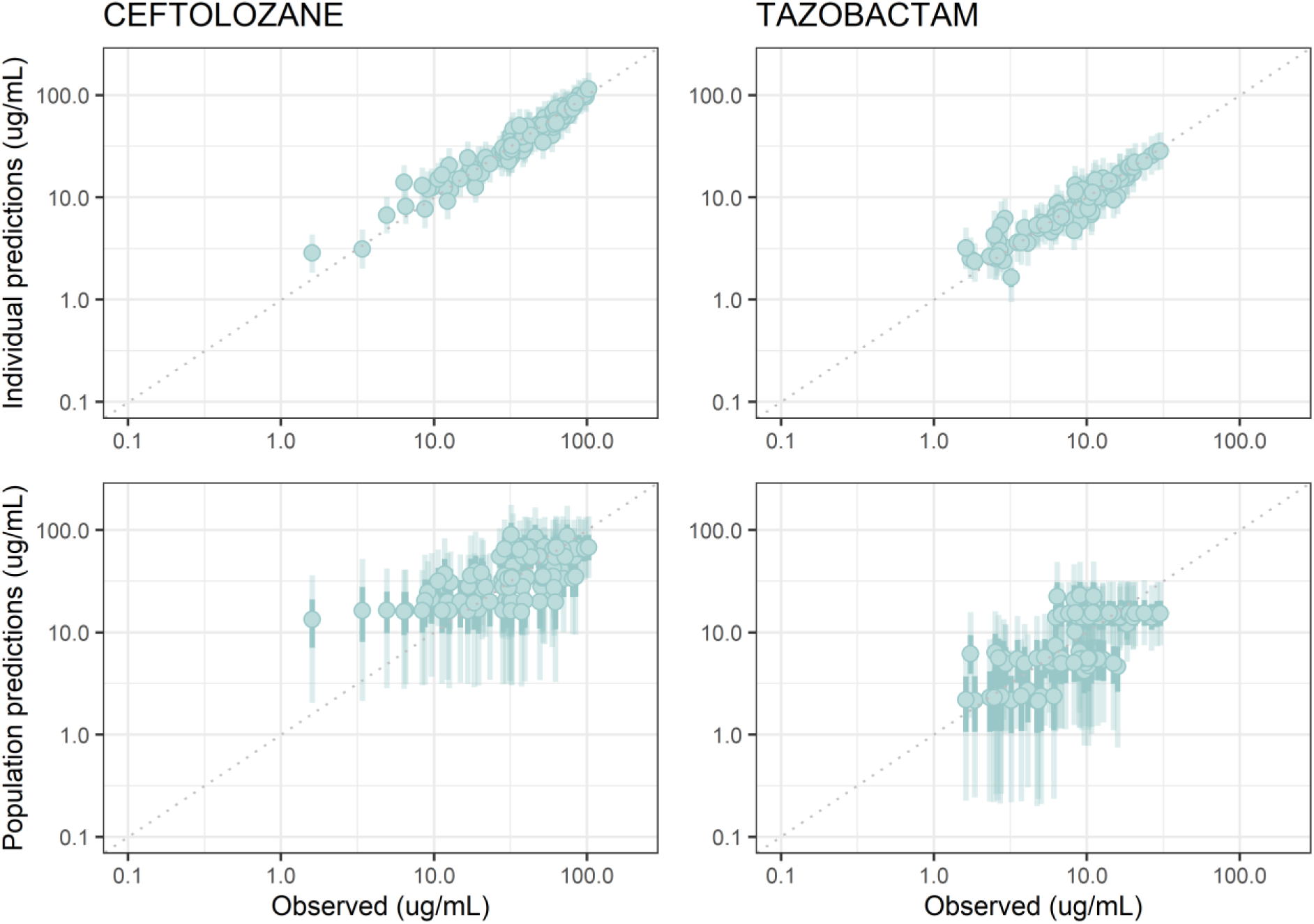
Posterior predictive intervals. Vertical bars show residual uncertainty; filled points indicate median.

The model’s performance was assessed via visual posterior predictive checks (VPCs), generated from 1000 simulated datasets using posterior parameters. The 10^th^, 50^th^, and 90^th^ percentiles were utilized to summarize both the observed data and the VPC predictions. Visual predictive checks (Figure 5) confirmed that the model accurately reproduced the observed data. For ceftolozane and tazobactam, the simulated median and 90% prediction intervals encompassed the majority of observed concentrations across dosing intervals, capturing both the multiphasic decline and repeated peak-trough pattern consistent with intermittent dosing (e.g., every 8 hours). Overall, the VPCs demonstrated good predictive coverage and distributional consistency between simulations and observations, validating the robustness of the Bayesian two-compartment model for simultaneous characterization of ceftolozane and tazobactam PK. The model thus provides a reliable quantitative framework for precision dosing and therapeutic drug monitoring in clinical settings. Individual posterior predictive checks for study participants are provided in the Supplementary Materials (Figures S7a and S7b).

**Figure 5.**
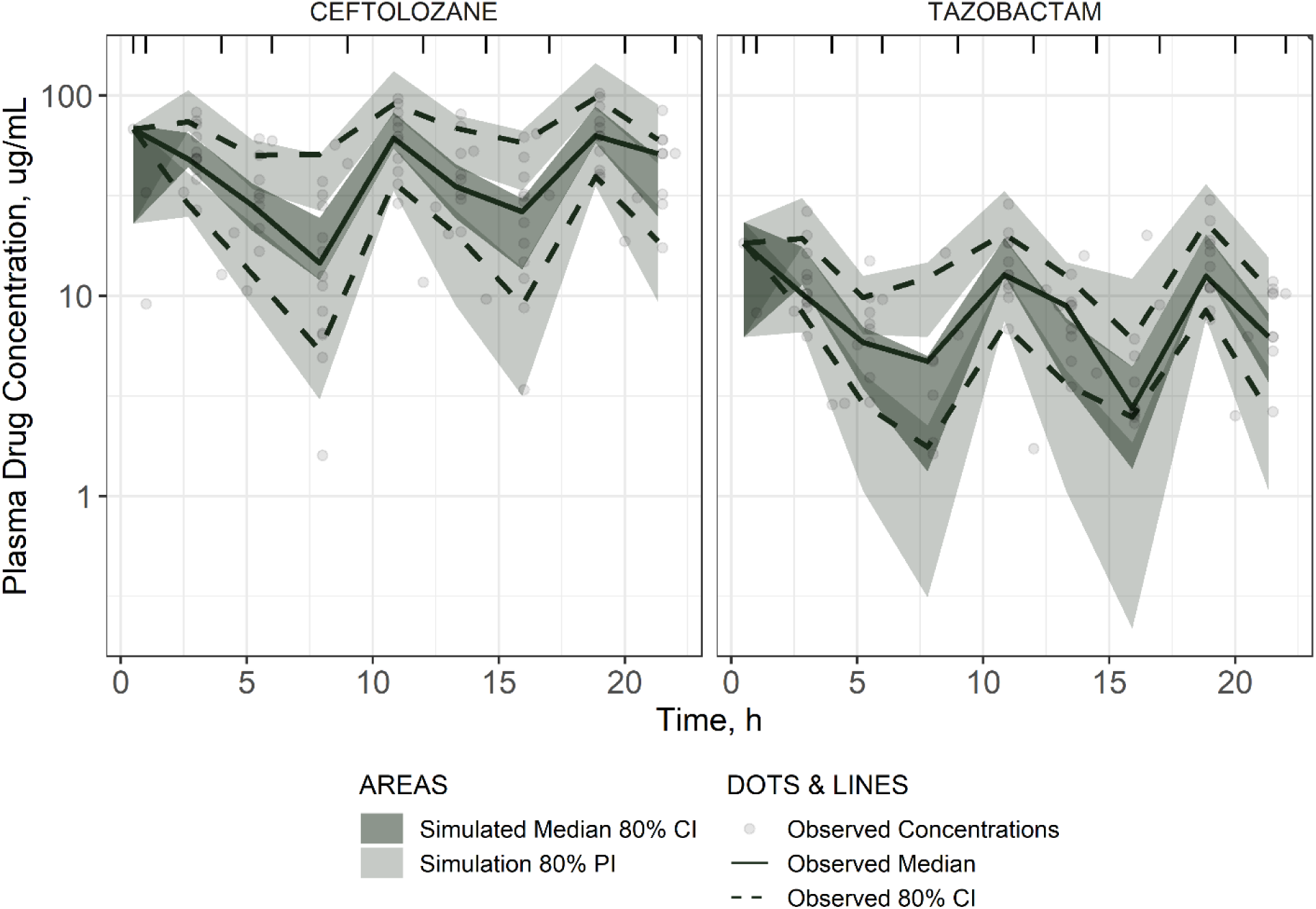
Visual posterior predictive checks comparing simulated posterior predictions with observed plasma concentrations of ceftolozane and tazobactam. Dashed lines represent 80% credible intervals for observed data, shaded areas indicate predictive intervals from simulations, and points correspond to measured concentrations.

### 4.4 Probability of Target Attainment (PTA) Analysis

Across all three simulation approaches, the PTA curves showed fairly similar patterns, maintaining values above 90% up to MICs of approximately 8-16 mg/L, followed by a sharp decline at higher MIC concentrations. The MIC values achieving 90% PTA were reasonably consistent, with medians (5^th^-95^th^ percentiles) of 21.86 mg/L (16.26-31.83 mg/L) for the one-compartment bootstrap, 20.04 mg/L (13.31-26.90 mg/L) for the one-compartment covariance approach, and 26.95 mg/L (21.79-31.07 mg/L) for the two-compartment Bayesian method. Using only the point estimate for the 1CMT covariance approach would produce a deterministic PTA curve without prediction intervals and a slightly higher MIC for 90% PTA (approximately 19-20 mg/L), leading to an overly optimistic assessment. In contrast, both uncertainty-inclusive methods, bootstrap (wider intervals due to non-parametric nature) and covariance sampling (narrower intervals assuming normality), provide more conservative and realistic estimates that account for parameter variability. The two-compartment Bayesian method yields less uncertain predictions by incorporating prior knowledge and produces more favorable PTA profiles, as ceftolozane and tazobactam concentrations remain above the target for a longer duration. It indicates that the 3 g q8h, 3-hour infusion regimen ensures robust probability of target attainment within the clinically relevant MIC range (Figures 6, S11). For the q12h regimen, all three simulations produced similar PTA profiles, though overall probabilities were lower compared to the q8h regimen at corresponding MICs. PTA remained above 90% up to approximately 4-8 mg/L, after which it declined more rapidly. The MIC values for achieving 90% PTA show reasonable similarity across methods, with medians (5^th^-95^th^ percentiles) of 19.62 mg/L (15.03-26.44 mg/L) for the one-compartment bootstrap method and 17.88 mg/L (12.41-22.94 mg/L) for the one-compartment covariance approach, and a median of 24.30 mg/L (20.23-27.71 mg/L) for the for the two-compartment Bayesian method. However, the reduced dosing frequency limits target attainment at higher MIC values (Figure 7, S12).

**Figure 6.**
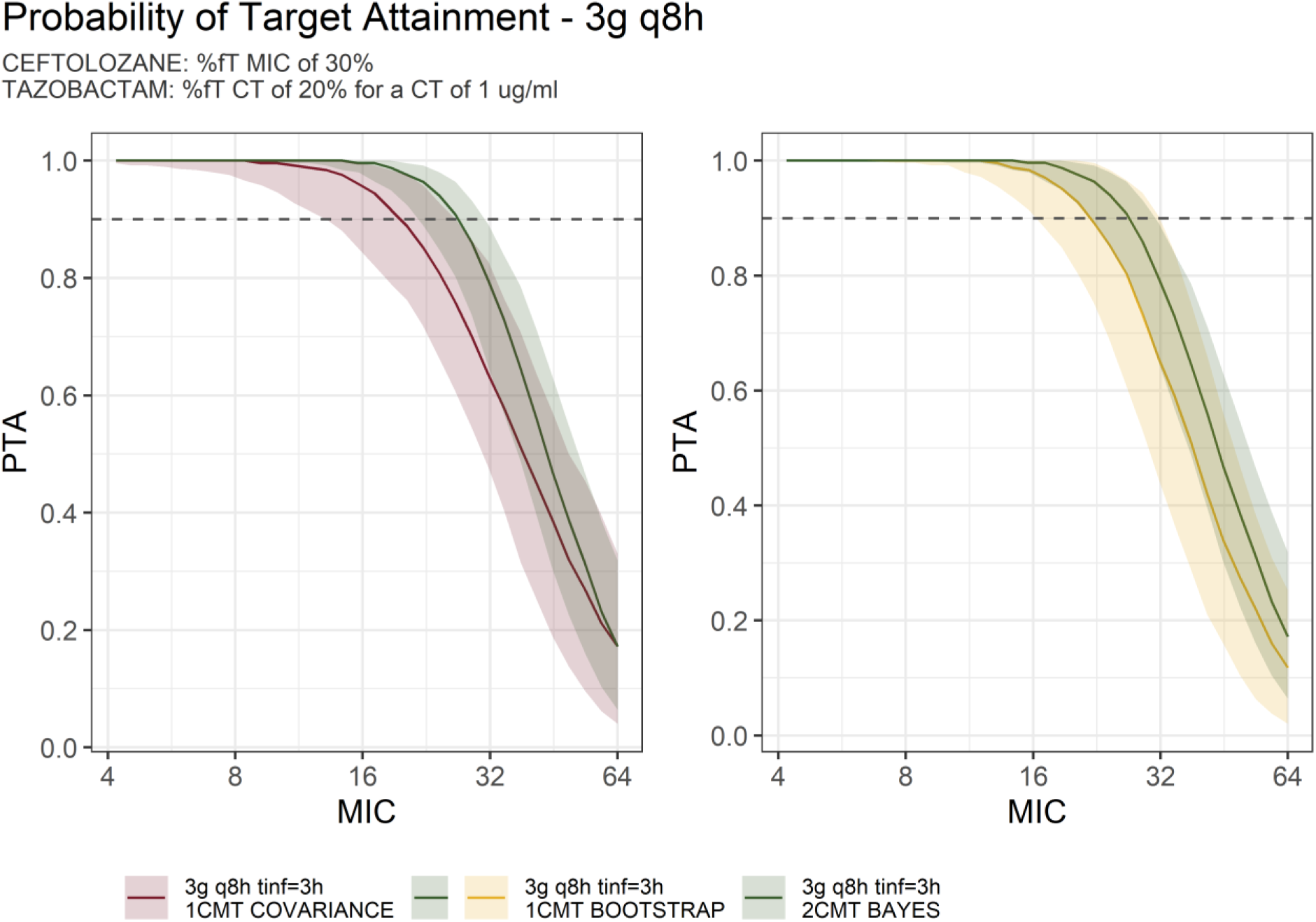
Probability of target attainment (PTA) for ceftolozane and tazobactam under the recommended 3 g q8h regimen with 3 h infusion. Median PTA values are shown as lines; shaded areas indicate 5–95% prediction intervals, and the dashed line denotes the 90% PTA threshold. Three simulation approaches are compared: Bayesian posterior predictions, one-compartment model (1CMT COVARIANCE), and 1CMT bootstrap-based simulations.

**Figure 7.**
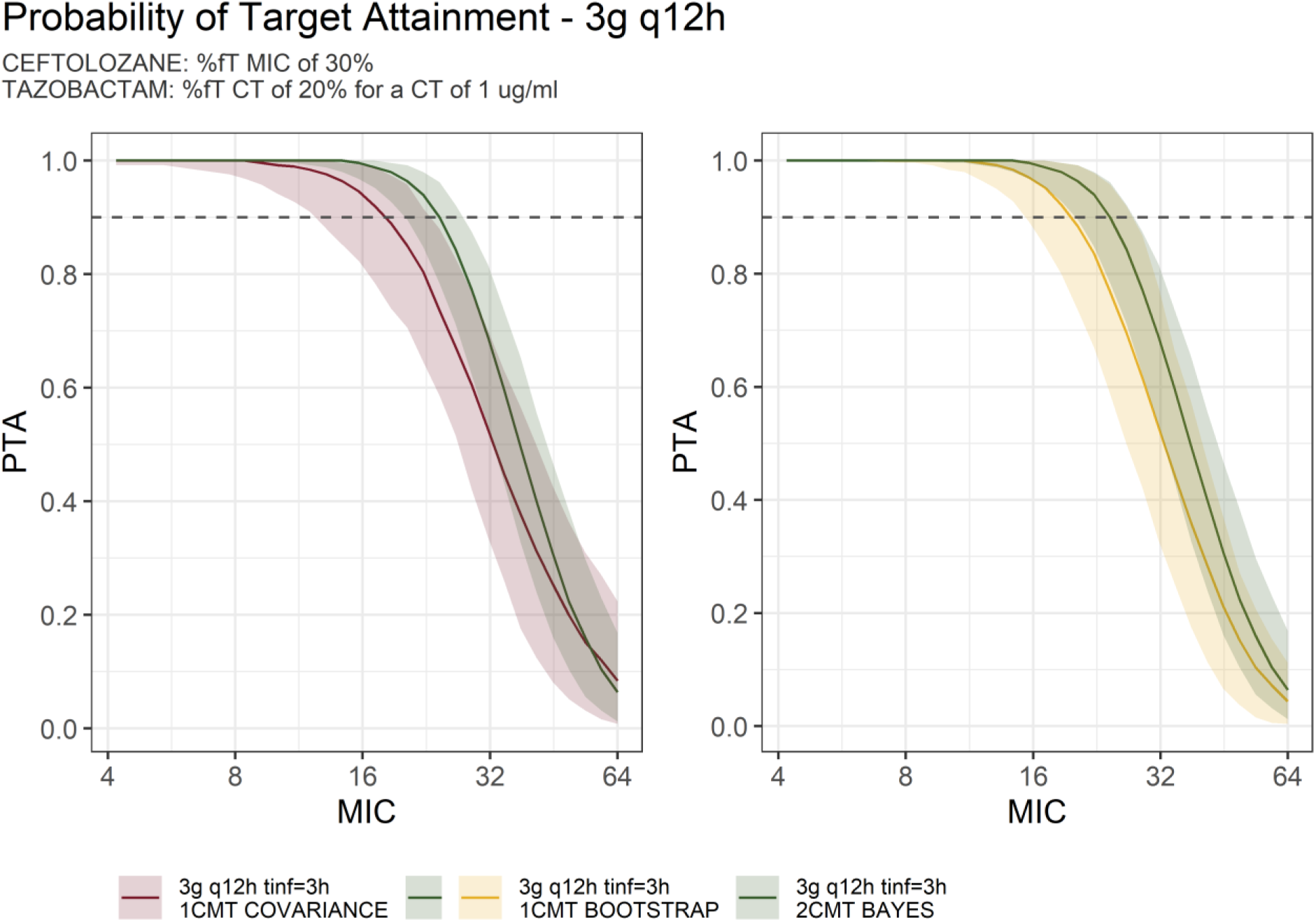
Probability of target attainment (PTA) for ceftolozane and tazobactam under a 3 g q12h regimen with 3 h infusion. Median PTA values and 5–95% prediction intervals are shown for Bayesian posterior predictions, one-compartment model (1CMT), and bootstrap-based simulations. This figure demonstrates model performance and PTA maintenance under an extended dosing interval

## 5. Conclusions

The primary objective of this study was to propose a population pharmacokinetic model for ceftolozane administered in combination with tazobactam in critically ill patients treated in intensive care units (ICUs). Antimicrobial therapy is a challenging field in pharmacology, primarily due to the need for precise dosing to achieve minimum inhibitory concentrations while minimizing the risk of antimicrobial resistance.

Using a one-compartment model, key parameters were determined: volume of distribution (*V1*) of 30.2 L and 55.5 L, and clearance (*CL*) of 6.37 L/h and 15.5 L/h for ceftolozane and tazobactam, respectively. Literature comparisons typically describe pharmacokinetics with a two-compartment model, which yielded the following estimates in the current analysis: central compartment volume (*V1*) of 25.3 L and 36.0 L, peripheral compartment volume (*V2*) of 45.0 L and 21.1 L, *CL* of 4.25 L/h and 15.9 L/h, and inter-compartmental clearance (*Q*) of 2.89 L/h and 103 L/h for ceftolozane and tazobactam, respectively. Model selection favored the one-compartment approach based on a lower objective function value (OFV: 820.5 vs. 877.8 for one-vs. two-compartment models). Moreover, two-compartment parameter estimates showed high uncertainty (relative standard error >30% for ceftolozane *V2*, tazobactam *V1* and *V2*, and tazobactam *Q*). If the true model is two-compartmental, the estimated parameters may be subject to systematic bias. The clearance (*CL*) parameter values for both drugs align with literature data. However, the estimated volume of distribution (*V1*) values in both the one-compartment model (30.2 L for ceftolozane and 55.5 L for tazobactam) and the two-compartment model substantially exceed those reported in other populations. Particular attention is warranted for the intercompartmental clearance (*Q*) of tazobactam, which was 103 L/h in our study, whereas literature values typically range much lower L/h (e.g., 3.13 L/h per Chandorkar et al., 2015 [27]; 4.06 L/h per Larson et al., 2019 [21]). This discrepancy was one reason for expanding the pharmacokinetic assessment of our study cohort.

In the Bayesian population PK framework, the prior and posterior distributions of key parameters: *CL, Q, V1,* and *V2,* demonstrated effective data-driven updating (Figure 3). For ceftolozane, the posterior distribution for *CL* was notably narrowed and shifted leftward (median approximately 5.74 L/h (5^th^-95^th^ percentile: 4.71-7.06) compared to *a prior* mode equal to 5.87 L/h (3.88-8.98), indicating slower systemic clearance than initially assumed. In contrast, *V1* showed a rightward shift (*posterior* median 22.33 L (17.69-27.00) versus *a priori* 10.66 L (7.10-15.86)), suggesting a larger central distribution. Comparable refinements were observed for tazobactam parameters, where the posterior for *CL* concentrated at 15.21 L/h (12.82-18.36) (from a broader *prior* spanning 20.73 L/h (13.71-30.98)), reflecting increased precision in estimating its elimination kinetics. Across all parameters posterior variances were reduced by approximately 50-80% (Figure 3), underscoring the informativeness of observed concentration-time data and the efficiency of Bayesian updating in constraining prior uncertainty derived from literature-based assumptions. Such posterior tightening enhances the precision of individual and population-level PK estimates, which is essential for dose individualization in antimicrobial therapy.

Building upon the robust predictive performance demonstrated by the PPCs, the Bayesian two-compartment model was extended to PD assessments via Monte Carlo simulations, evaluating the probability of target attainment (PTA) under various dosing regimens. This integration of PK modeling with PD endpoints highlights the model’s clinical relevance, particularly in scenarios with limited patient data or variable dosing conditions.

Under the q8h and q12h dosing regimens with a 3-hour infusion, all three simulation approaches: one-compartment model (1CMT), bootstrap-based simulations and Bayesian posterior predictions, produced consistent PTA profiles. Median PTA values exceeded 90% up to MICs of approximately 8-16 mg/L across all methods. The 1CMT model generated slightly lower PTA values at higher MICs due to its simplified structure, while bootstrap simulations showed wider 5-95% prediction intervals, reflecting greater uncertainty. When patient data is limited, the Bayesian model provides a distinct advantage. By integrating prior pharmacokinetic information with sparse individual data, it enables stable and individualized posterior predictions even in small cohorts, maintaining realistic estimates of variability. In contrast, the one compartment model may underestimate variability, and bootstrap analysis becomes less reliable with few observations.

The results of this study enhance the prediction of plasma concentrations of ceftolozane and tazobactam in critically ill patients, thereby improving the overall efficacy of this antibiotic combination. By refining a Bayesian two-compartment pharmacokinetic model, the findings facilitate optimized dosing regimens for intravenous administration. This approach, validated through visual predictive checks and probability of target attainment under standard dosing with infusion, supports precision medicine in antimicrobial therapy, particularly for resistant infections, while minimizing adverse effects associated with under- or overdosing.

## 6. Data and Code Availability

The R code, data, and Stan code used to analyze the data are publicly available in the GitHub repository https://github.com/pokunska/ceftolozane-tazobactam.git

## 7. Acknowledgments

This research received no specific grant from any funding agency in the public, commercial, or not-for-profit sectors. Authors’ Contributions: PMO, PW and MC contributed to writing; PMO and PW, and MC to conceptualization and methodology; GR to analytical method development and assay; PMO to data curation; PW and PMO to statistical methodology, data analysis, and visualization; and MB, ER, PP, MS, MC to clinical care of patients and data collection.

